# Investigating Facial Expression Processing during Fast Periodic Visual Stimulation using Highly Variable Stimuli

**DOI:** 10.1101/2025.02.14.638300

**Authors:** David Vandenheever, Haleigh Davidson, Jennifer Kemp, Zack Murphy, Autumn Kujawa, Catherine Shi, Michael Nadorff, Kayla Bates-Brantley, MacKenzie Sidwell

## Abstract

Facial expression recognition is a fundamental aspect of human social interaction, enabling effective communication and emotional understanding. Fast Periodic Visual Stimulation (FPVS) paradigms have recently emerged as a powerful approach for studying facial expression processing. However, previous studies often utilized identical base stimuli, making it difficult to disentangle neural responses to low-level perceptual differences from those reflecting conceptual discrimination of emotion. By introducing variability in our stimuli, we aimed to overcome these limitations and investigate neural responses to facial expressions of anger, fear, happiness, and sadness. Using EEG, robust oddball responses were observed across participants at both individual and group levels, demonstrating the paradigm’s sensitivity even with brief recordings and limited post-processing. Significant neural responses were detected across key regions of interest, with the right occipito-temporal region showing the strongest activity, consistent with its role in high-level facial expression processing. This study highlights the effectiveness of the FPVS paradigm for examining emotional processing using naturalistic stimuli and provides a framework for future research into neural mechanisms underlying facial emotion recognition in diverse and pathological populations.

## 1. Introduction

Recognizing facial expressions is a critical component of human social interaction, enabling us to infer the internal emotional states of others. This ability allows for rapid and adaptive responses to social cues, which are essential for effective communication and survival in group settings (Poncet, et al., 2019). From an evolutionary perspective, the ability to process, identify, and discriminate facial expressions quickly is crucial, particularly in scenarios where members of a group may display contrasting emotions, such as anger or fear, signaling aggression or the need for caution, respectively (Schettino, et al., 2020). Humans have developed complex neural mechanisms to categorize facial expressions and generalize across varying facial identities, a process that involves both rapid visual discrimination and higher-order conceptual judgments (Lau & Dzhelyova, 2020).

Scientific investigations into facial expression recognition have frequently employed scalp electroencephalography (EEG) due to its high temporal resolution and ability to measure direct brain activity in various populations (Poncet, et al., 2019). Traditional event-related potential (ERP) studies have identified early neural responses to expressive faces, typically within the first 150 ms post-stimulus, suggesting rapid attentional processing (e.g., (Eimer & Holmes, 2002), (Batty & Taylor, 2003), (Dzhelyova, et al., 2017)). However, these findings are often confounded by uncontrolled low-level image properties, such as luminance or contrast differences, which may contribute to observed effects (Dzhelyova, et al., 2017). Consequently, the precise neural mechanisms underlying facial expression recognition remain elusive, necessitating novel approaches to address these methodological challenges (Dzhelyova, et al., 2017) (Coll, et al., 2019) (Rossion, 2014).

Fast Periodic Visual Stimulation (FPVS) paradigms have recently emerged as a powerful alternative for studying facial expression processing. FPVS leverages the presentation of stimuli at a strict periodic rate, eliciting neural responses at predefined frequencies in the EEG spectrum. This approach provides an objective, high signal-to-noise ratio (SNR) measure that is both implicit and efficient, isolating category-selective processes without the need for post-hoc subtraction (Rossion, 2014) (Quek & Rossion, 2017). Notably, FPVS is used to investigate implicit facial processing and is argued to discriminate faces at the conceptual level, including identity and emotion, as evidenced by its modulation or absence of responses for inverted faces (Coll, et al., 2019) (Dzhelyova, et al., 2017).

Several studies have successfully employed FPVS to explore facial expression processing. Dzhelyova et al. (2017) demonstrated robust oddball responses to emotional expressions such as fear, disgust, and happiness, highlighting distinct neural signatures over occipito-temporal regions. Similarly, Poncet et al. (2019) identified expression-specific EEG responses to a range of emotions, revealing partially distinct neural substrates for different facial expressions. Luo and Dzhelyova (2020) further investigated pairwise contrasts of expressions, observing qualitative differences in neural responses, particularly for happiness and fear. Naumann et al. (2025) extended FPVS research to children, demonstrating robust expression-change responses, with variations in frequency power linked to expression intensity.

While these studies underscore the utility of FPVS, they also reveal certain methodological limitations. All these studies used a single facial identity per condition trial, introducing variability between successive images by changing image sizes. However, as Coll et al. (2019) noted, this approach can lead to FPVS discrimination based on low-level visual properties, such as differences in contrast or luminance (e.g., a change in contrast due to the teeth being shown in the oddball, but not the base, stimulus), rather than the emotional content of the oddball images. Supporting this, Schettino et al. (2020) reported oddball responses even for inverted faces, further suggesting that low-level features can confound FPVS responses.

Coll et al. (2019) attempted to address some of these challenges by incorporating 20 different facial identities (10 male), reducing the likelihood of low-level confounds. However, they converted the images to grayscale, adjusted for differences in luminosity, cropped the images around the natural line of the face thereby removing other features such as hair and shoulders, and they presented the images against a white background. The artificial standardization of low-level stimulus properties is believed to degrade stimulus quality and ecological validity which also negatively affects generalization (Quek & Rossion, 2017) (Rossion, et al., 2015) (Retter, et al., 2021). Furthermore, De Rosa et al. (2022) showed that FPVS responses can be evoked by the relative token frequency of base and oddball stimuli. This means that differences in how frequently specific images or identities are presented between the base and oddball conditions can also elicit an oddball response. This underscores the need for even more variety in facial identities beyond the 20 used by Coll et al. (2019).

The use of full-color images and greater variability is further supported by Rossion et al. (2015), who emphasized the strength of using large, highly variable stimulus sets in terms of low-level properties. Color images add another level of variability and are also highly diagnostic for face detection, with behavioral evidence indicating that colored faces are detected faster than grayscale ones (Rossion, et al., 2015). Building on this foundation, the present study introduces several methodological improvements to enhance the ecological validity and robustness of FPVS paradigms. We considerably increase the number of facial identities while presenting the stimuli in full color, uncropped, and include natural backgrounds, hair, and shoulders, ensuring greater stimulus variability and addressing the potential for token frequency effects.

By addressing these limitations, this study aims to provide a more comprehensive and ecologically valid assessment of facial expression processing. The use of highly variable, naturalistic stimuli allows for the investigation of expression discrimination in a manner that generalizes across identities and contexts, offering new insights into the neural mechanisms underlying this critical social ability.

## 2. Methods

### 2.1 Participants

Forty-one adults (aged 18-28, mean age 20.8, 21 females) participated in the study. They were recruited from the Mississippi State University student population. All participants had normal or corrected-to-normal vision and hearing. Ethical approval for all procedures were obtained from the Mississippi State University Institutional Review Board. All participants provided written informed consent before participating and were free to withdraw from the study at any time. Three datasets were removed due to technical difficulties (for two datasets the EEG data was not recorded) or excessive noise (one dataset). Therefore, results are reported for 38 participants.

### 2.2 Stimuli

The face images used were selected from the FACES database, which has been widely validated and recognized (Ebner, et al., 2010). The FACES database includes 2,052 pictures of faces from 171 individuals aged between 19 to 80 (85 females). There are two images per person and expression (six facial expressions). We only used five facial expressions (neutral, angry, fear, sad, and happy). We resized all images from 819 × 1,024 pixel resolution to 158 × 197 (w x h) pixels, 96 dpi. The images subtended 5° x 6.25° visual angle (w x h) and were presented on a 24” monitor with 60 Hz refresh rate.

### 2.3 Procedure

Participants were seated 80 cm from the monitor. During the FPVS paradigms, face images were presented on the monitor for 83 ms, followed by an 83 ms blank interval before the next image appeared. Images were presented in sequences consisting of five images. The first four images were selected from the base category, while every fifth image was drawn from the oddball category. This design induces two distinct steady state responses: one at the base presentation frequency of 6 Hz and another at the oddball frequency of 1.2 Hz. Refer to Figure 1 below for a visual representation of the paradigm. Participants were required to detect changes in the color of a fixation cross displayed on the screen throughout each trial. The cross changed from blue to red for a duration of 400 ms, occurring between six to eight times at irregular intervals within each trial. This setup ensures that participants maintain their focus on the screen without engaging with the stimuli. We used four FPVS conditions, each lasting about 122 seconds and presented in random order. Each condition had neutral face images as the base stimuli with happy, sad, fearful, and angry faces as oddball stimuli, respectively.

**Figure 1.**
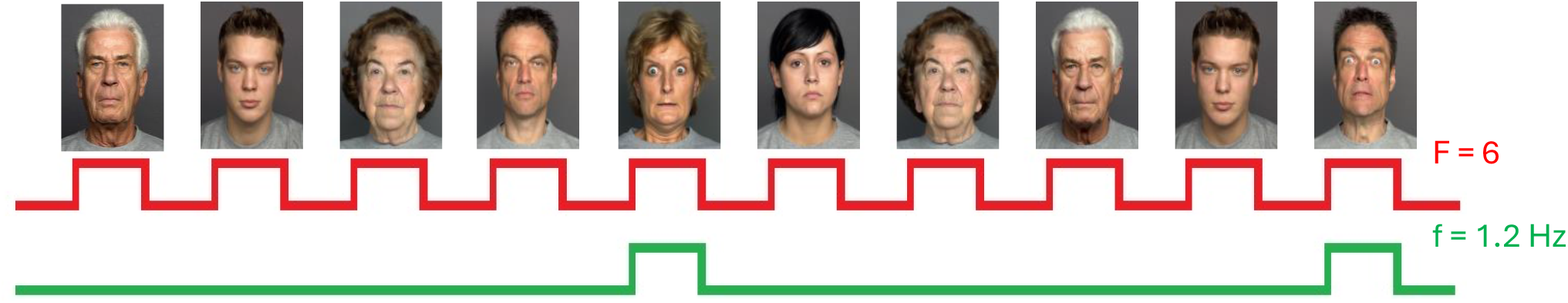
Schematic illustration of the experimental paradigm. Stimuli are presented at a rate of 6 Hz. In each 122 s stimulation sequence, base stimuli are selected from a large pool with oddball images presented at fixed intervals of one every fifth stimuli or 1.2 Hz. The example in this figure corresponds to condition C, fear faces. Note that only six faces are shown due to copyright and use agreement.

### 2.4 EEG recording

We recorded EEG from 64 active scalp electrodes using a BioSemi ActiveTwo system (BioSemi, Amsterdam, Netherlands). EEG signals were sampled at 256 Hz from 64 channels with reference set to the average of right and left mastoids.

### 2.5 EEG processing and analysis

ALL EEG data processing was performed using Matlab (Mathworks Inc.) and the EEGLab toolbox. Data was segmented to include 2 s before and 3 s after each sequence, resulting in 127 s segments. Data was digitally bandpass filtered between 0.1 to 100 Hz using a finite impulse response (FIR) filter. Bad channels were identified (exceeding 5 standard deviations from mean kurtosis value) and replaced using spherical interpolation of the neighboring channels. On average less than 3% of channels were interpolated per participant. Finally, data was re-referenced to a common average of all electrodes. Notably, we didn’t perform artefact rejection as the FPVS method is assumed to be relatively immune to artefacts and meaningful responses can be obtained without correction for blinks or other artefacts (Rossion, 2014). This is due to the fact that the neural response of interest in FPVS is confined to a very specific frequency. This simplifies the analysis procedure, making it highly reproducible across studies.

The preprocessed data segments were cropped to an integer number of 1.2 Hz cycles, beginning about 2 s after the onset of the sequence until approximately 120 s. The first 2 s of each segment were excluded to avoid any contamination by the initial onset response. A fast Fourier transform was then applied to these segments and normalized amplitude spectra were extracted for all channels and participants (Dzhelyova, et al., 2017). Previous research has shown a robust SSVEP response to the oddball frequency and many of its harmonics with oddball detection more reliably and accurately measured when including the harmonics of the oddball response (Retter, et al., 2021) (Stothart, et al., 2020). To select the statistically significant harmonics to include in the estimation of response amplitude, we expressed the amplitude at each harmonic frequency as a *z*-score relative to the mean and standard deviation of the amplitude of the 20 neighboring bins of the FFT (10 bins on each side, excluding the immediately adjacent and two most extreme values (Dzhelyova, et al., 2017) (Naumann, et al., 2025). A *z*-score greater than 1.64 (i.e., *p* < 0.05, one-tailed) was the criterion to consider the amplitude at that frequency significantly greater than the amplitude at neighbouring frequency bins (Dzhelyova, et al., 2017) (Rossion, et al., 2015) (Quek & Rossion, 2017) (Lochy, et al., 2015). We investigated the number of significant harmonics, pooled across all electrodes and on group-level, starting at 1.2 Hz and moving upward. We also investigated significant harmonics at specific regions of interest (ROIs): occipital (channels Oz, O1, O2), frontal (Fz, FCs, AFz), and lateral occipito-temporal (left: PO7, P7, P9, right: PO8, P8, P10). Once a harmonic failed to reach significance, we identified the range for that condition as the last harmonic that was significant. Harmonics that related to the standard frequency (e.g. 6 Hz, 12 Hz…) were excluded (Stothart, et al., 2017).

To increase the signal to noise ratio, previous studies have either used more but shorter (35-60 s) trials per condition, or segmented their trials into shorter segments, and then average the trial in the time-domain before computing the FFT (Coll, et al., 2019) (Dzhelyova, et al., 2017) (Dzhelyova, et al., 2020) (Poncet, et al., 2019). We investigated the effect of this on our z-scores by segmenting our trials into three epochs of approximately 40 s, averaging in the time-domain, and then performing the FFT. We then computed the z-scores as outlined above.

To quantify the facial expression discrimination response at the oddball frequencies, we calculated baseline-corrected amplitudes (BCA) on individual subjects’ spectra by subtracting the mean amplitude of the 20 surrounding bins (10 bins on each side, excluding the immediately adjacent and two most extreme values) and summed for the first 20 harmonics, excluding the harmonics corresponding to the base response and its harmonics (6 Hz, 12 Hz etc.). The z-scores confirmed that responses were significant up to 17 harmonics for some conditions, and even up to 20 harmonics for some individuals. Including nonsignificant harmonic responses should not have a great influence on the summed BCA since the BCA should equal zero or at least be close to zero if there is no discernible response at a specific oddball harmonic (i.e. it should look similar to the mean of the neighboring amplitudes which is subtracted). The responses were quantified over all channels as well as the previously defined ROIs. Paired two-sided t-tests were used to do comparisons between the different facial expression conditions (Lutz, et al., 2024).

Lastly, we also calculated SNR values at each frequency of interest expressed as the amplitude value divided by the average amplitude of the 20 surrounding frequency bins (10 on each side, excluding the immediately adjacent and two most extreme values). SNR spectra were used for data visualization. Grand-averages of the SNR and BCA were computed for each condition and electrode separately.

## 3. Results

The analysis revealed robust responses to facial expression changes at the oddball frequency (1.2 Hz) and its harmonics, as indicated by the high signal-to-noise ratio (SNR) values across all conditions (Figure 2). All four facial expressions (angry, fearful, happy, and sad) elicited significant mean z-scores above the threshold (z > 1.64) at the oddball frequency and its harmonics, pooled across all electrodes and at specific ROIs (Table 1). Across conditions, the highest mean z-scores were observed in occipital (O) and right occipito-temporal (rOT) regions, with more participants demonstrating significant z-scores in these regions compared to frontal (F) or left occipito-temporal (lOT) areas. Notably, the 120-second trials consistently showed more participants with significant responses, while the 40-second trials revealed a higher number of significant harmonics, especially for sad and happy faces, with up to 17 harmonics reaching statistical significance. The number of participants with significant responses varied slightly by ROI and facial expression. The occipital and right occipito-temporal regions consistently showed the highest number of participants with significant z-scores, underscoring the role of these areas in facial expression discrimination.

**Table 1.** Grand-average z-scores at oddball frequency and its harmonics for the different ROIs. Also shown are the number of significant harmonics and the number of participants that had significant mean z-scores up to the significant harmonics. ROIs: All = all 64 electrodes; O = Occipital; lOT = left Occipito-temporal; rOT = right Occipito-temporal; F = Frontal.

| Condition | ROI | One 120 s trial |  |  | Three 40 s trials averaged |  |  |
| --- | --- | --- | --- | --- | --- | --- | --- |
|  |  | Significant Harmonic | Mean z-score | Significant Participants | Significant Harmonic | Mean z-score | Significant Participants |
| Angry | All | 13 | 2.17 | 27 | 13 | 2.11 | 25 |
|  | O | 13 | 4.61 | 33 | 13 | 4.46 | 33 |
|  | lOT | 13 | 2.35 | 29 | 13 | 2.29 | 29 |
|  | rOT | 13 | 3.09 | 33 | 13 | 3.07 | 31 |
|  | F | 13 | 2.10 | 26 | 13 | 2.00 | 23 |
| Fear | All | 13 | 2.22 | 28 | 14 | 2.14 | 27 |
|  | O | 13 | 4.66 | 32 | 14 | 4.54 | 33 |
|  | lOT | 13 | 2.47 | 31 | 14 | 2.31 | 32 |
|  | rOT | 13 | 3.24 | 35 | 14 | 3.11 | 34 |
|  | F | 13 | 2.27 | 26 | 14 | 2.18 | 24 |
| Happy | All | 14 | 2.23 | 30 | 17 | 2.02 | 26 |
|  | O | 14 | 4.78 | 34 | 17 | 4.12 | 34 |
|  | lOT | 14 | 2.48 | 32 | 17 | 2.29 | 31 |
|  | rOT | 14 | 3.06 | 35 | 17 | 2.74 | 34 |
|  | F | 14 | 2.29 | 28 | 17 | 1.99 | 22 |
| Sad | All | 14 | 2.10 | 27 | 17 | 1.95 | 24 |
|  | O | 14 | 4.45 | 33 | 17 | 4.03 | 32 |
|  | lOT | 14 | 2.21 | 29 | 17 | 2.11 | 27 |
|  | rOT | 14 | 2.95 | 34 | 17 | 2.70 | 33 |
|  | F | 14 | 2.07 | 24 | 17 | 1.93 | 22 |

**Figure 2.**
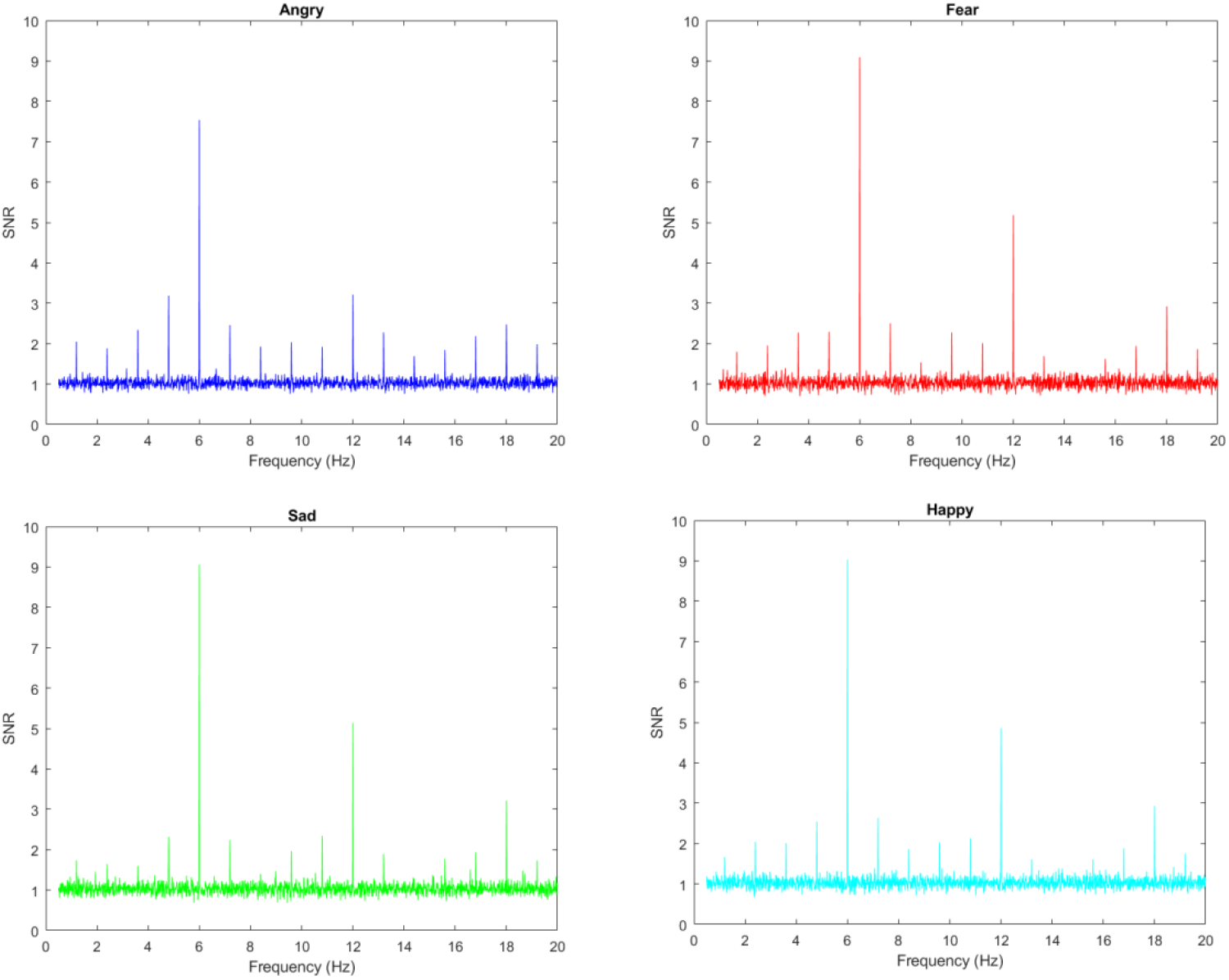
Grand-average SNR for the oddball responses over Right Occipital-temporal electrodes. The high SNR responses at the base rate (6 Hz) and its harmonics (12 Hz and 18 Hz) are clearly visible, as are the oddball response (1.2 Hz) and its harmonics.

Analysis of BCA showed no significant differences between facial expressions in frontal and occipital ROIs. However, significant differences emerged in the left and right occipito-temporal regions, where sad faces elicited lower responses compared to angry, fearful, and happy faces (Figure 3).

**Figure 3.**
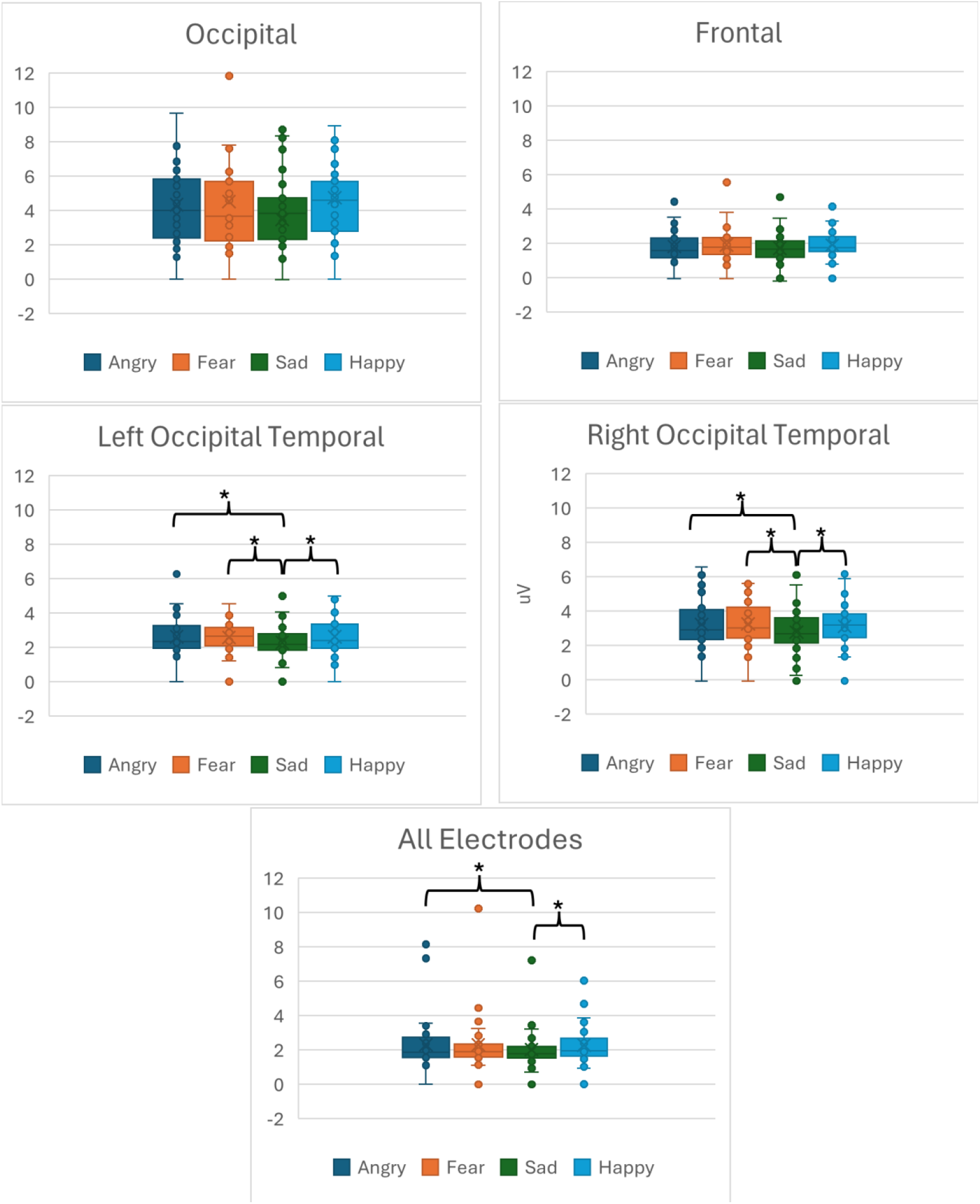
Grand-average summed baseline-corrected oddball amplitudes for facial expression changes at the five ROIs. A response above 0 reflects signal > noise. Statistically significant differences (p < 0.05) are indicated by the *.

These results demonstrate the effectiveness of the FPVS paradigm in eliciting robust neural responses to facial expressions, with specific regional and participant-level variations that highlight the nuanced processing of different emotions.

## 4. Discussion

Our study demonstrates robust responses to facial expression changes at both individual and group levels, even with a single two-minute recording, highly variable stimuli, and no artefact removal in post-processing. This supports the feasibility of using FPVS paradigms with naturalistic, variable image sets for facial emotion processing, addressing limitations identified in prior studies that relied on low-variability or standardized stimuli. Prior approaches often struggled with interpreting oddball responses, as facial expressions can be discriminated on various levels, including low-level visual features, face-specific processing, and emotional-semantic recognition (Coll, et al., 2019). By introducing greater variability into the stimuli, our study reduces the risk of confounding factors and ensures that responses reflect higher-level processes rather than simple low-level differences (De Keyser, et al., 2018). Additionally, using diverse, full-color stimuli enhances ecological validity and eliminates the need for strict standardization of features like spatial frequency, luminance, and contrast, which can otherwise bias results (Coll, et al., 2019) (De Keyser, et al., 2018).

The findings align with previous studies showing maximal face-selective activity over right occipito-temporal (rOT) regions, further confirming their role in high-level facial expression and emotion processing (Dzhelyova, et al., 2017) (Rossion, 2014) (Xu, et al., 2017). Despite including frontal ROIs to account for emotion-related processes (Schmidt, et al., 2010) (Liu, et al., 2020), we found no significant differences between expressions in these regions. However, significant z-scores were observed in frontal areas, suggesting that these regions are indeed associated with broader emotion processing or may reflect the recruitment of higher-level areas involved in integrating emotional and contextual information. This suggests that while frontal asymmetry is associated with broader emotional processing, occipito-temporal areas may dominate in discriminating specific facial expressions.

Interestingly, sad faces elicited lower responses compared to the other expressions, consistent with findings by Leleu et al. (2018). This could reflect differences in how sadness is processed at neural and behavioral levels. Other studies have also reported increased responses for positive stimuli, referred to as a positivity bias (Schettino, et al., 2019). Furthermore, fear and angry faces might carry greater survival relevance, as they convey critical information for threat detection and response. This evolutionary significance could explain a bias toward heightened responses for these expressions compared to sadness. However, we did not observe significant differences between fear and happiness, contrasting with studies such as Poncet et al. (2019) and Luo and Dzhelyova (2020), which reported distinct neural signatures for these emotions. Methodological differences, including our use of highly variable stimuli, may account for these discrepancies.

While our study provides strong support for the utility of FPVS paradigms, it is not without limitations. First, although we employed a highly variable and ecologically valid set of stimuli, the generalizability of our findings to other paradigms or stimuli remains to be explored. Although our stimuli had more variability, all the faces were still from the same vantage point and presented against a uniform background. It would be interesting to investigate emotion processing using this paradigm but with even more variability in facial stimuli, such as differences in angle and orientation. Furthermore, while our sample size was relatively large for FPVS studies, the diversity of our participants was limited as it consisted primarily of college students. Future studies should aim to include a more diverse cohort to enhance the generalizability of the findings. Additionally, this paradigm holds promise for exploring facial emotion processing in pathological populations, such as individuals with autism spectrum disorder (ASD), as demonstrated by van der Donck et al. (2019), who used FPVS to investigate facial emotion sensitivity in children with ASD. Expanding the application of this technique to alternative or additional pathological populations, such as individuals with anxiety or depression (i.e., internalizing disorders), could provide valuable insights into potential differences in emotion processing and the underlying neural mechanisms.

In conclusion, our results validate the FPVS approach for studying facial expression processing using ecologically valid, variable stimuli. By demonstrating robust neural responses across diverse conditions and regions, this study advances the field’s understanding of emotion-specific processing and highlights the utility of FPVS for individual-level assessments. Most importantly, we found robust oddball responses at the individual level in short recording times (only 2 minutes per condition) with minimal post-processing and no artifact removal. This efficiency and reliability support the potential of FPVS as a clinical screening tool. These findings underscore the need for further research employing the FPVS approach in clinical populations to explore its diagnostic and prognostic capabilities, potentially paving the way for its integration into clinical neuropsychological assessments.

## Notes

### Competing Interest Statement

The authors have declared no competing interest.

